# Spatial differentiation of proteome in cervical cancer tissues using Imaging Mass Spectrometry

**DOI:** 10.1101/2025.04.12.648495

**Authors:** Aastha Baliyan, Ipsita Mohapatra, Sayanti Paul, Abass Alao Safiriyu, Santosh Kumar Mondal, Deepak Mishra, Amit Kumar Mandal

## Abstract

Cervical cancer, which is the fourth most common gynaecological cancer across the globe, has a poorly understood molecular pathogenesis and etiology. Current methods of diagnosis are based on cytology, histology and presence of Human Papillomavirus (HPV). Shortcomings of these methods lie with poor quality of smears which is very common in Papanicolaou (pap) smears, limited sensitivity in terms of early detection, dependence on presence of HPV, etc. Due to its high sensitivity and non-targeted approach, Matrix Assisted Laser Desorption Ionization based Imaging Mass Spectrometry (MALDI-IMS) might be advantageous to understand the pathogenesis, molecular mechanism, precise identification of surgical margins and identification of novel biomarkers. Since it can also identify proteins in the extracellular matrix, it is especially beneficial for the tissue types with sparse cells and excessive extracellular matrix. Although tissue proteome profiling for cervical cancer were reported, the heterogeneous distribution of proteins across cervical cancer tissues haven’t been explored. In this study, we employed a non-targeted MALDI-IMS based approach to profile the spatial distribution of proteins within cervical cancer tissues. We observed overexpression of Keratin 5 and Prelamin A/C in the region of cervical cancer tissues which were categorically labelled with cancerous morphology using histopathological examination. Both these proteins have been earlier associated with progression and aggressiveness of other cancers like breast and prostate cancers. However, no such reports are available for cervical cancer. Further studies are required on a large dataset to validate and quantitate these proteins as biomarkers for early diagnosis and prognosis of cervical cancer.

## INTRODUCTION

Cervical cancer is the fourth most common gynaecological cancer across the world, leading to approximately 660,000 new cases and 350,000 deaths annually [1]. It has the highest prevalence and mortality rates in low- and middle-income countries where India shares 19% of the global burden of cervical cancer cases [2,3]. The survival rate of cervical cancer in India is 80% within 1 year of diagnosis which drops to 39% within 5 years of diagnosis.

One of the known causes of cervical cancer is Human Papilloma Virus (HPV). HPV are sexually transmitted, non-enveloped, DNA viruses of the family papillomaviridae [4]. Although the majority of cervical cancers are caused by HPV, approximately 5.5 - 11% of cervical cancers are HPV-negative and have a poorly understood etiology [5]. There is a lack of biomarkers to identify such HPV-negative cases which may escape detection and treatment, leading to poor prognosis [6]. Although HPV infection is primary requisite, it is not sufficient to develop cervical cancer [7]. The details of molecular mechanisms governing the molecular pathogenesis of cervical cancer is still largely unknown [8]. In general, the carcinoma cervix originates in the transformation zone. The cervical intraepithelial neoplasia (CIN) lesions are graded based on the degree of dysplastic cells where CIN1 is mostly benign cellular manifestation involving one-third of the epithelium; CIN2 is moderate dysplasia up to two thirds of the epithelium, and CIN3 is severe dysplasia including full thickness of the epithelium and is considered a direct precursor to cancer [9,10].

Current screening and early detection of cervical cancer is either by detecting the presence of HPV, liquid cytology or Papanicolaou smear (Pap smears) [11,12]. The gold standard for diagnosing cervical cancer in a tissue is histopathology based analysis. All of these methods have their own sets of limitations like assuming that all cervical cancer cases are HPV-positive, relying on the human expertise, low overall sensitivity/specificity and relatively high variability [13-17]. Cervical cancer continues to present challenges in early stage diagnosis, thus warranting a critical need to develop robust biomarkers that can unambiguously discriminate between cancerous and non-cancerous tissues regardless of the HPV status of the tissue. Since there are a number of gaps in our knowledge of cervical cancer pathogenesis, detection and treatment, therefore, it is crucial to study the molecular profile of cervical cancer tissues in detail in order to identify novel biomarkers suitable for early diagnosis and better prognosis.

Mass spectrometry is a versatile tool that is widely employed for proteomics profiling and biomarker development for various diseases. It can provide a tool for unbiased label-free identification and quantitation of hundreds of proteins simultaneously. However, classical proteomics is suitable for analysis of serum samples or tissue lysates which end up erasing the information regarding the localization and differential expression profile of different proteins. MALDI-IMS provides molecular detail of the tissues, providing spatial localization of protein biomarkers within heterogeneous tissue environments which can help to define the surgical margins with better precision [18-20]. Although there are numerous reports of employing MALDI-IMS to study cancerous tissues, till date, there is no report on analysis of cervical cancer tissues using MALDI-IMS platform. In this study, we employed MALDI mass spectrometry based tissue imaging to analyze the cervical cancer tissues in order to visualize the heterogeneous distribution of various proteins across the cervical cancer tissues and to identify potential protein biomarkers associated with the cervical cancer.

## MATERIALS AND METHODS

### Materials

Very high purity (>99.9%) TPCK treated trypsin, β-octyl glucopyranoside, ammonium bicarbonate, Acetonitrile (ACN) (LCMS grade), Water (LCMS grade), Formic acid (FA) and α-cyano hydroxycinnamic acid (CHCA) were purchased from sigma. Analytical grade ethanol, chloroform, acetic acid, xylene, lithium carbonate, eosin, hematoxylin, D.P.X. mountant for microscopy and potassium sulfate were purchased from SRL. LC-MS grade water and trifluoroacetic acid (TFA) were purchased from Thermofisher Scientific. The Superfrost™ slides were purchased from fisher scientific. Rapigest and yeast enolase digestion standard was purchased from Waters,UK. Sunchrom sprayer was used to spray trypsin and matrix on the tissues and the images were recorded on Synapt G2-Si (Waters, UK) with MALDI source.

### Ethics statement

The cervical cancer tissues were obtained from All India Institute of Medical Sciences, Kalyani, India (AIIMS Kalyani) after obtaining written consent from the participants. The study was approved by the Institutional Ethical Committee, AIIMS Kalyani (IEC approval no. IEC/AIIMS/Kalyani/Meeting/2022/05). All the procedures were carried out in accordance with the available guidelines and regulations for human participants in research.

### Selection of subjects

Patients presenting with cervical cancer who visited the O.P.D. of Department of Obstetrics and Gynaecology at AIIMS, Kalyani, were invited to participate in the study under the supervision of the gynecologist. The details of the study were explained to the patients following which the written consent was obtained and the sample was collected.

### Sample acquisition and cryosectioning

Six punch biopsy samples were collected from patients who were suspected of cervical cancer, after obtaining written consent and snap-frozen in liquid nitrogen. The tissue samples were stored at -80 °C until further use. The tissues were thawed to -20 °C in a cryostat and subsequently mounted on a microtome chuck with minimal amount of mounting media. Tissues were fixed such that the exposed surface is not in contact with the mounting media. Tissue sections of 8 - 10 µm thickness were thaw mounted on charged slides. The sectioned slides were stored at -80 °C until further use. The tissue sections were first imaged using MALDI-IMS and were subsequently processed for Hematoxylin and Eosin (H and E) staining.

### MALDI Imaging

#### Preparing slides for MALDI Imaging

For fixing the tissues and removing the lipids and salts, the slides were treated using Carnoy’s method as follows: 70% ethanol (30 seconds), 100% ethanol (30 seconds), Carnoy’s solution containing 60% chloroform, 30% ethanol and 10% acetic acid (120 seconds), 100% ethanol (30 seconds), 0.2% aqueous TFA (30 seconds) and 100% ethanol (30 seconds). Following this, the slides were vacuum dried for 30 minutes to remove all the solvents. The slides were placed in a box containing saturated aqueous potassium sulfate solution at the bottom such that the slides are not in contact with the solution. The box containing the slides was then incubated at 90 °C for 5 minutes to heat denature the proteins [21]. The slides were vacuum dried for 15 minutes to remove excess moisture. The slide was then sprayed with 0.1 mg/ml trypsin solution prepared in 50 mM ammonium bicarbonate buffer (pH 7.4) containing 0.1% β-octyl glucopyranoside using Suncollect sprayer/spotter, at a rate of 7 µL/minute for 10 layers. The slide was then dessicated for 5 minutes to dry excess liquid to prevent delocalization of proteins. The slide was then incubated in a humid chamber containing saturated potassium sulfate solution at 37 °C overnight. After digestion, the slide was vacuum dried for 45 minutes and sprayed with 5 mg/ml CHCA solution prepared in 50/50 ACN/0.8% aqueous TFA, using Suncollect sprayer/spotter at flow rate of 10 µL/minute for 2 layers, 15 µL/minute for 6 layers and 20 uL/minute for 2 layers. The slides were then dried for 60 minutes before imaging in MALDI qToF mass spectrometer (Synapt G_2_-Si, Waters, UK).

#### MALDI based peptide imaging

The slides sprayed with CHCA were scanned and uploaded into the HDI™ software to define the areas to be imaged. The slides were loaded into the MALDI mass spectrometer Synapt G_2_-S_i_ (Waters, UK) which was calibrated using Red Phosphorus in range 1000 – 5000 m/z. The spectra were acquired in positive polarity in Sensitivity mode with a resolution of 20,000. The Trap Collision Energy (Trap CE) was set to 2 V and Transfer CE was set to 10 V. The pixel size of the image was set to 50 microns in both X and Y coordinates. Laser repetition rate was 1000 with an energy of 480. The quad profile during the acquisition was set to Auto, for a mass range of 1000 – 4500 m/z.

#### Imaging Data analysis

The acquired data was processed using HDI™ software using trypsin autolysis peaks at m/z 2163.07 as internal lockmass with a tolerance of 0.25 amu and intensity threshold of 250 counts. Top 1000 peaks were selected to generate images and the m/z window was set to 0.02 m/z. The intensities of all peaks were normalized by TIC (Total Ion Count) where the intensities of all peaks are divided by the sum of absolute intensities of all peaks. Molecular images were created by the software using a cartesian coordinate system for a given mass-to-charge (m/z) ratio. Data points in the x and y coordinates represent the distribution of proteins across the tissue section and the signal intensity (i) correlates with the abundance of the corresponding protein molecule. The m/z signals which showed a clear image of distribution within the tissue were selected as proteins of interest. These proteins were then identified and characterized using a nanoLC-MS/MS based proteomics platform using commercial facility SAIF at IIT-Bombay.

### Histopathology

After MALDI imaging, the tissues were subjected to H and E staining using standard protocol and imaged at 20X resolution. In brief, to stain the tissues, they were treated with methanol bath for removing the matrix, followed by sequential treatment with with 95% ethanol (30 seconds), 70% ethanol (30 seconds), 50% ethanol (30 seconds), distilled water (10 minutes) to rehydrate the tissue, Harris hematoxylin (10 minutes), 0.05% lithium carbonate (60 seconds), distilled water (10 seconds, twice), Eosin stain (30 seconds) and distilled water (10 seconds each, thrice). Following the staining, the tissue was then dehydrated by treating with ethanol gradation as follows: 50% ethanol (30 seconds), 70% ethanol (30 seconds), 90% ethanol (30 seconds), 100% ethanol (60 seconds) and xylene (5 minutes). On this tissue the coverslip was mounted using D.P.X mounting media and imaged at 20X magnification. These slides were read by the pathologist to mark the distribution of cancerous cells within the tissue.

### Proteomics

#### Sample preparation

Two milligrams of tissue was dried in a lyophilizer for 2 hours and frozen in liquid nitrogen before crushing to powder using a micropestle. The crushed powder was suspended in a lysis buffer containing 10 mM Tris (pH 7.4), 150 mM NaCl, protease inhibitor, 1mM EDTA and 1% β-octyl glucopyranoside. The suspension was sonicated for 30 minutes at 35% amplitude with intermittent cooling (10 seconds on cycle, 30 seconds off cycle). The lysate was then incubated at 4 °C with occasional mixing by gently inverting the tubes. The lysate was centrifuged at 12,000 rpm for 30 minutes, at 4 °C to remove the cell debris. Protein concentration in the supernatant was measured using Bradford assay. Samples were then denatured by adding 0.2% Rapigest and incubating at 80 °C for 15 minutes, followed by reduction (using 5 mM DTT, 60 °C, 30 minutes) and alkylation (10 mM IAM, RT, 1 hour). The proteins were then digested by trypsin (1:10 enzyme:protein molar ratio) by incubating overnight at 37 °C. After digestion, rapigest was inactivated by acidifying by addition of 0.5% Formic acid (v/v) and incubating at 90 °C for 90 minutes.The samples were then centrifuged at 12,000 rpm for 30 minutes at 4 °C. Supernatant was collected and spiked with internal standard (yeast enolase digestion standard 50 fmoles/µL).

#### nanoLC-ESI-MS/MS based fractionation

For tissue proteomics, the samples were commercially analyzed at SAIF facility, IIT Bombay. The spiked samples were analyzed on the EASY-nanoLC UHPLC system (Thermo Scientific, Germany). Briefly, 600 ng of total tryptic digest of the experimental sample was loaded directly into a C18 column PepMap RSLC (2 µm, 100Å x 50 cm). The aqueous phase consisted of 0.1% aqueous FA (solvent A) and the organic phase consisted of 85:15 (ACN:water) + 0.1% FA (solvent B). The flow rate of solvents through the column was 300 nL/min. A 60 min run program was designed, starting with a linear gradient of solvent B from 5-20% in 20 min, followed by 20-50% of solvent B in 10 min, and finally 50-95% of solvent B in 8 min. The eluted peptides were analyzed in a mass spectrometer with an orbitrap mass analyser (Q Exactive Plus Biopharma, Thermo Scientific, San Jose, California, USA).

#### Data acquisition

The data were acquired on the Thermo Scientific Xcalibur software (Version 4.2.28.14) in the mass range of 350−1600 m/z at 70000 FWHM resolution. The instrument was operated in positive ion mode with a source temperature of 350 °C, capillary voltage of 2.0 kV. After a survey scan, tandem MS was performed on the most abundant precursors exhibiting charge states from 2 to 7 with the intensity greater than 5.0e3, by isolating them in the quadrupole. The automatic gain control (AGC) target for MS/MS was set to 2.0e5 and the maximum injection time limited to 120 msec for each microscan.

#### Proteomics data analysis

The acquired data was processed and analyzed using the Thermo Proteome Discoverer software version 2.2 against the *homo sapiens* database from uniprot.

### Correlating the histopathology data with molecular images and proteomics data

The H and E images were overlapped with the molecular images obtained from the mass spectrometer using the HDI software. The peptides which showed differential expression across samples were identified by performing proteomics analysis using nanoLC-MS/MS. The identified proteins were analyzed using pathway and functional analysis (Table 1).

**Table 1:**
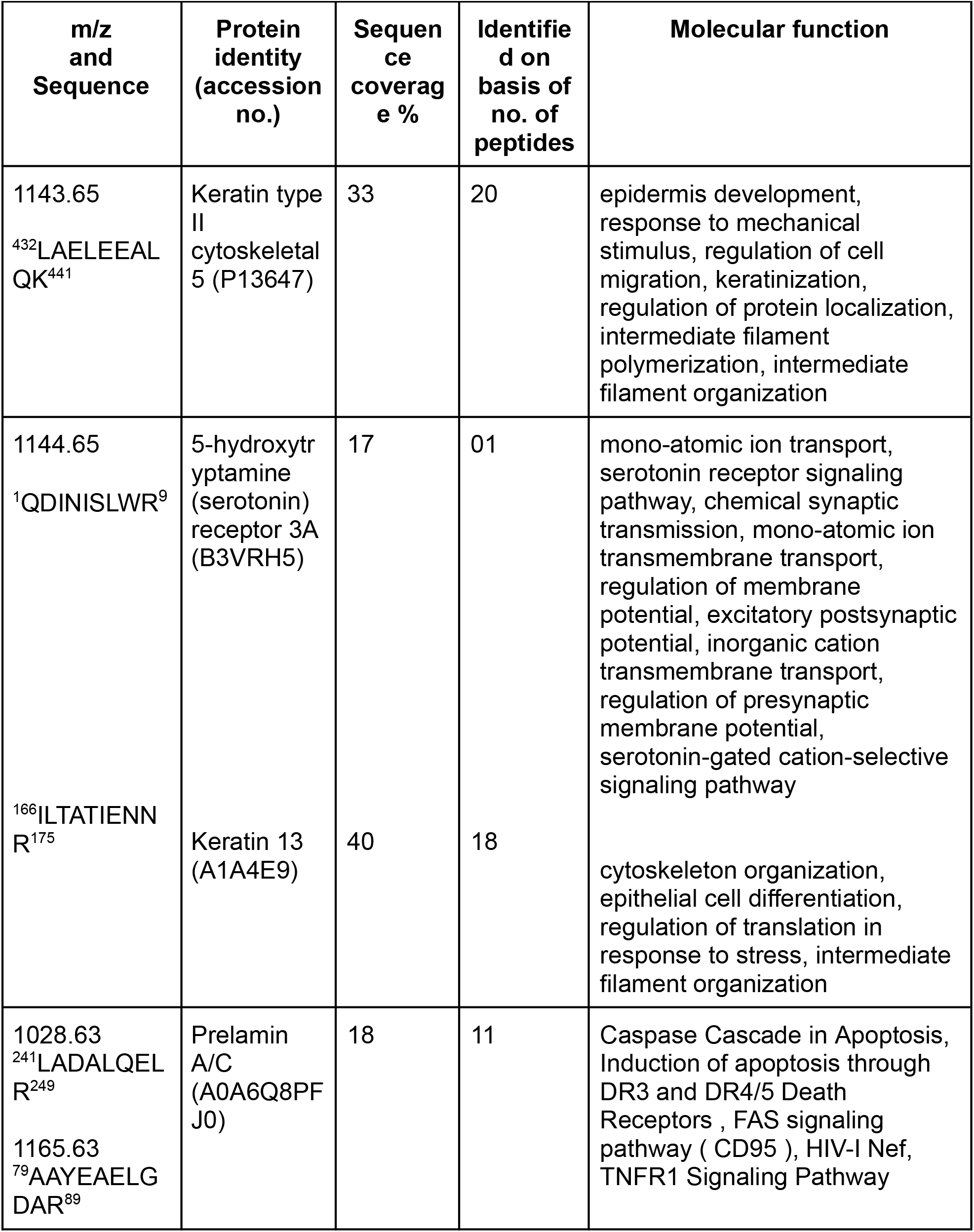
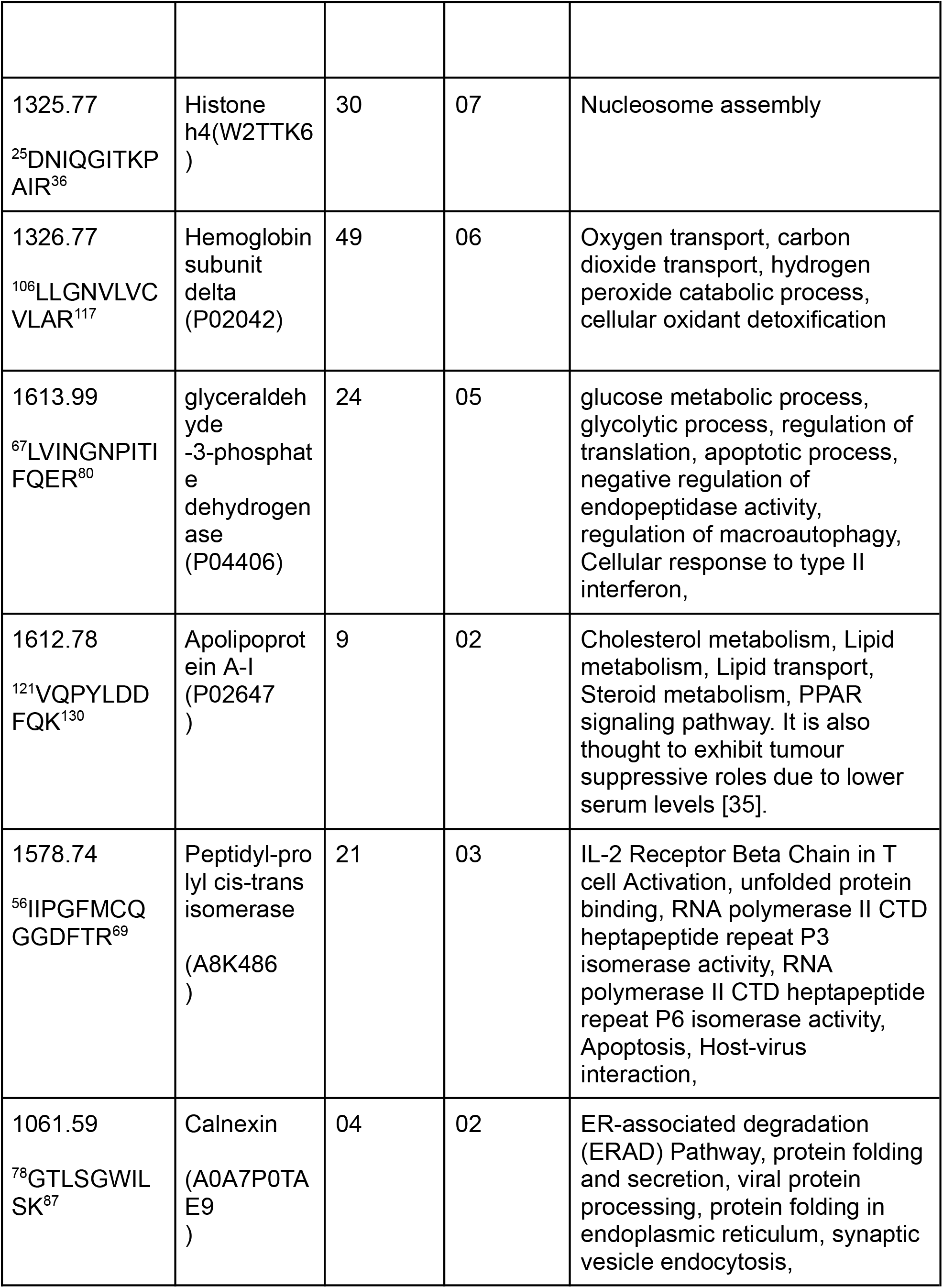
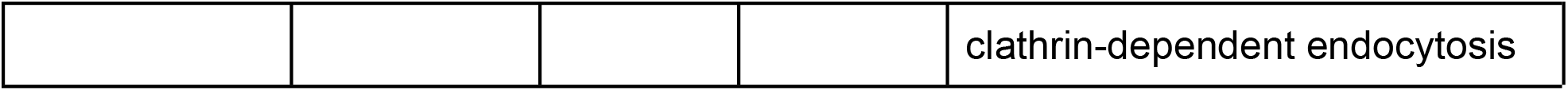
Proteomics data identified 10 proteins which are involved in various pathways in normal cellular homeostasis.

## RESULTS AND DISCUSSION

Current diagnostic and screening methods for cervical cancer involve mainly pap smear test (cytology), HPV testing, and histopathological analysis by a trained pathologist. Pap smear and HPV genome based tests rely on the assumption that all precancerous lesions are HPV positive and can be tested for oncogenicity by presence of HPV genome. While this is true in the majority of cases, a small but significant subset (5.5 - 11%) of cervical cancer cases can be HPV-negative [5]. Such HPV-negative cases can be difficult to detect through molecular screening alone. Hence, the diagnosis is entirely based on cytology and histopathology, which might lead to false negative diagnosis especially in early stages of the disease [17]. In the histopathological study, the tissues are characterized based on the morphological features of the cells, tissue architecture and nuclear staining. This is particularly troublesome in cervical cancer cases, because of poor quality of the pap smears, cellular sparsity in biopsy samples, and subjective nature of diagnosis. Therefore, it is important to have a robust, sensitive and highly specific diagnostic method to rule out both false positive and negative diagnosis.

Over the past couple decades, mass spectrometry based proteomics and tissue imaging have emerged as powerful techniques in cancer research, biomarker discovery and pathway analysis of disease pathology. MALDI-IMS based tissue imaging can provide diagnostic insights from cellular contents as well as the extracellular matrix and microenvironment of the tissue, and therefore, is not negatively affected by the cellular sparsity like conventional cytological or histopathological methods. However, its application to cervical cancer, in the context of biopsy-based research, still remains limited. A comprehensive database search on ProteomeXchange with the filters for “Homo sapiens” and “cervical cancer” yielded only 36 results, out of which, only 6 studies were focused on biopsy samples from patients [22]. This highlights the need for further proteomic studies of cervical cancer samples to map the molecular portrait of cervical cancer pathogenesis. MALDI-IMS provides a unique advantage by enabling the visualization of cancer boundaries and heterogeneity, which is not possible in conventional in-solution proteomics experiments. However, employing MALDI-IMS for cervical cancer is particularly challenging due to the unsuitability of cryosectioning these tissues. This might be due to high water content, cellular heterogeneity leading to uneven freezing rates, low structural rigidity, cellular sparsity due to more extracellular matrix, etc.

In the current study, we employed a MALDI-IMS based tissue imaging platform to image various proteins across six cervical cancer tissue sections and thereby, to identify protein patterns that are exclusively overexpressed in cancerous regions of the tissue. All those tissues were confirmed to be cancerous through histopathological examination. Figure 1 represents histological images (H and E) images of two representative cervical cancer tissue sections. Figure 1a illustrates a tissue (hereby referred as sample A) entirely filled with cancerous cells, where the insets 1 and 2 show the clusters of cells exhibiting high nuclear-to-cytoplasmic (N:C) ratio. Figure 1b illustrates a tissue (hereby referred to as sample B) having some portions of cancerous cells (inset 1) while others are healthy cells (inset 2). Figures 2 and 3 represent the molecular images of proteins generated from MALDI-IMS imaging of sample A and B, respectively. Sample A was diagnosed as cancerous based on histopathological examination, whereas sample B was diagnosed as negative based on pap smear test but H and E staining revealed cancerous regions within the tissue. The molecular images of different proteins were observed at varying concentrations throughout the tissues which were formed by representing the mass spectral relative intensities as heat maps. For each m/z value in the mass spectra, an intensity plot or image is created where the heat map of ion intensity in relation to a color scale shows the variation in intensity of each protein throughout the tissue section. In these intensity heatmaps, yellow represents the highest relative intensity, followed by red, blue and black as minimum intensity. Figure 4 shows the comparative analysis of the three protein images alongside the H and E images of the respective tissue sections from sample A (top panel) and B (bottom panel). In order to identify and characterize the proteins observed in the molecular images, in-solution proteomics was performed on a chunk of tissue sample. Various peptides from the imaging experiments were identified using the proteomics approach as explained in materials and methods. Proteomics data was analyzed using the Proteome Discoverer software against Homo sapiens database. The results were then filtered by removing the proteins showing low confidence based on scores, peptides (less than 2 peptides per protein) and sequence coverage (Table 1). The molecular functions of the identified proteins were mapped using the DAVID database [23]. We identified two differentially expressed proteins (KRT5 and Prelamin A/C) through the representative tryptic peptides. KRT5, imaged through tryptic fragment 1143.6 m/z (^432^LAELEEALQK^441^), and Prelamin A/C, imaged through tryptic fragment 1165.6 m/z (^79^AAYEAELGDAR^89^), are differentially expressed in cancerous tissues as compared to a non-specific tryptic fragment imaged at 1030.4 m/z. Figure 5 represents the overlaid molecular images of KRT5 and Prelamin A/C obtained for sample B with the H and E images of the same tissue section. H and E based tissue imaging is the gold standard in defining cancerous tissues. Overlaid images of KRT5 and Prelamin A/C with the histopathological analysis clearly indicates that these two proteins are overexpressed in the cancerous regions within the tissue section.

**Figure 1:**
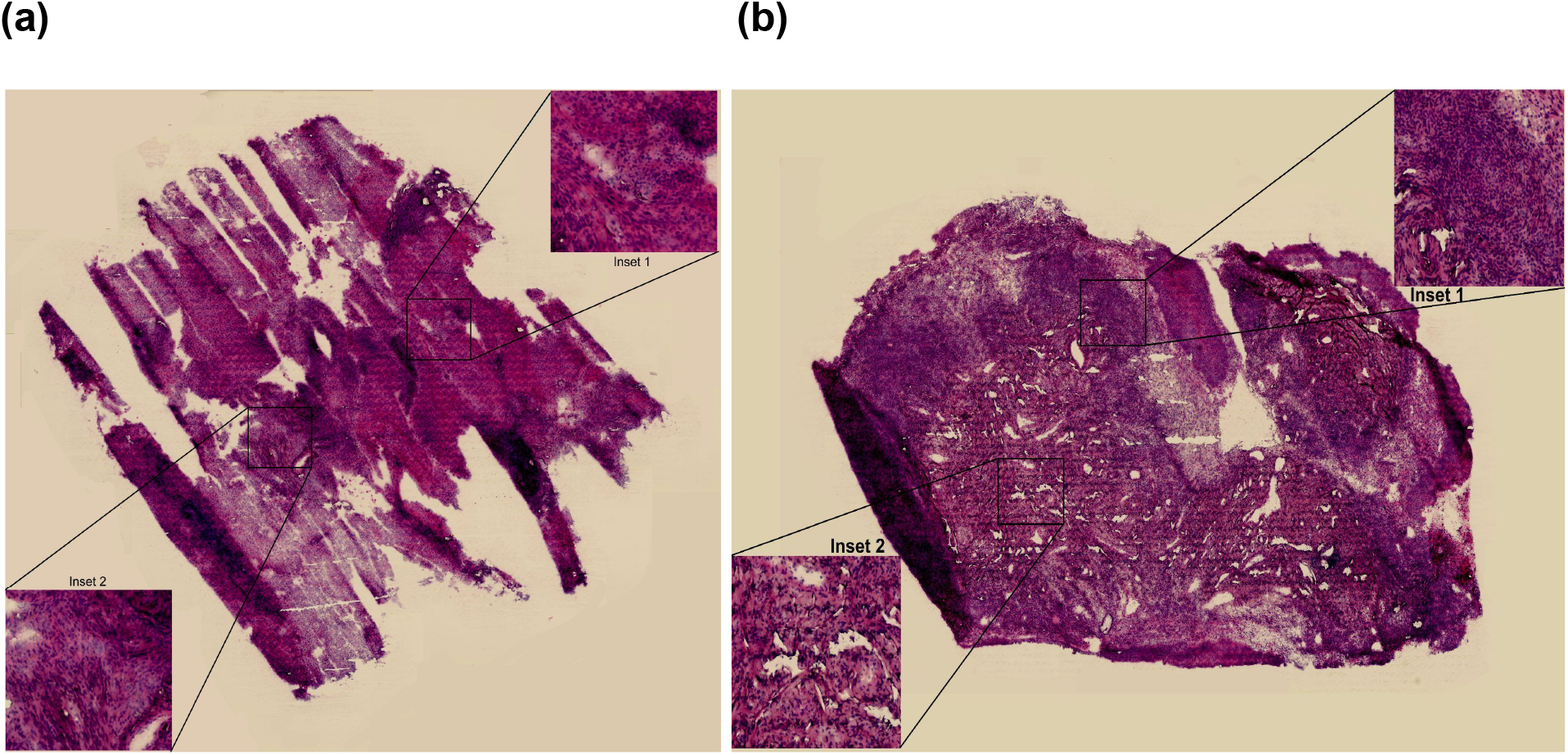
Histopathological images of H and E stained cervical cancer representative tissues. (a) Squamous cell carcinoma, wherein the entire tissue is cancerous and is devoid of any healthy cells. (b) A tissue partially cancerous (inset 1) and partially healthy (inset 2). The pap smear of this tissue was negative and hence was diagnosed to be negative for malignancy based on cytology.

**Figure 2:**
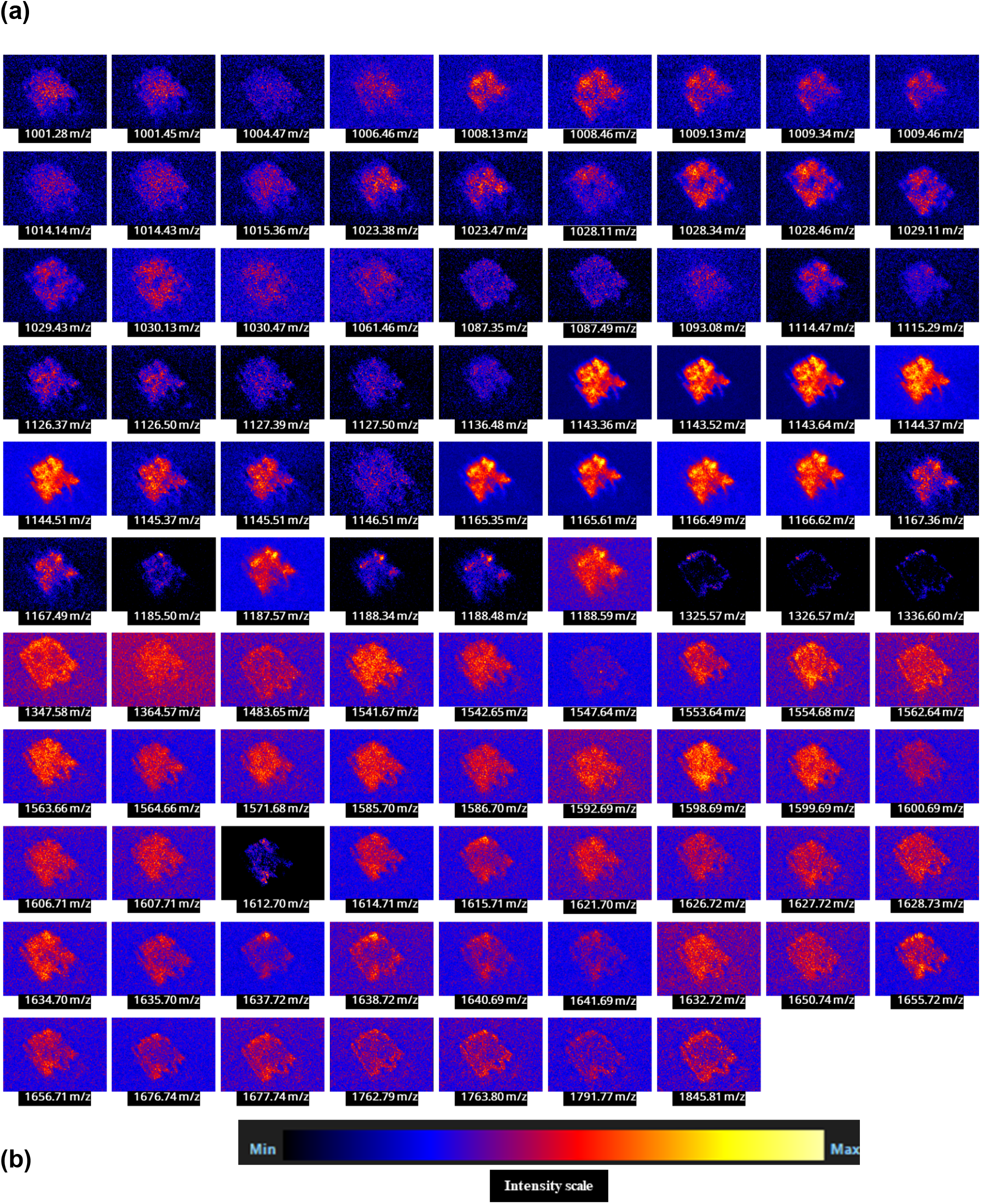
(a) The molecular images of different peptides as represented by their respective m/z values in sample A. (b) Intensity scale of the heatmaps where black represents the minimum relative intensity followed by blue, red and yellow representing the highest relative intensity of an ion across the tissue.

**Figure 3:**
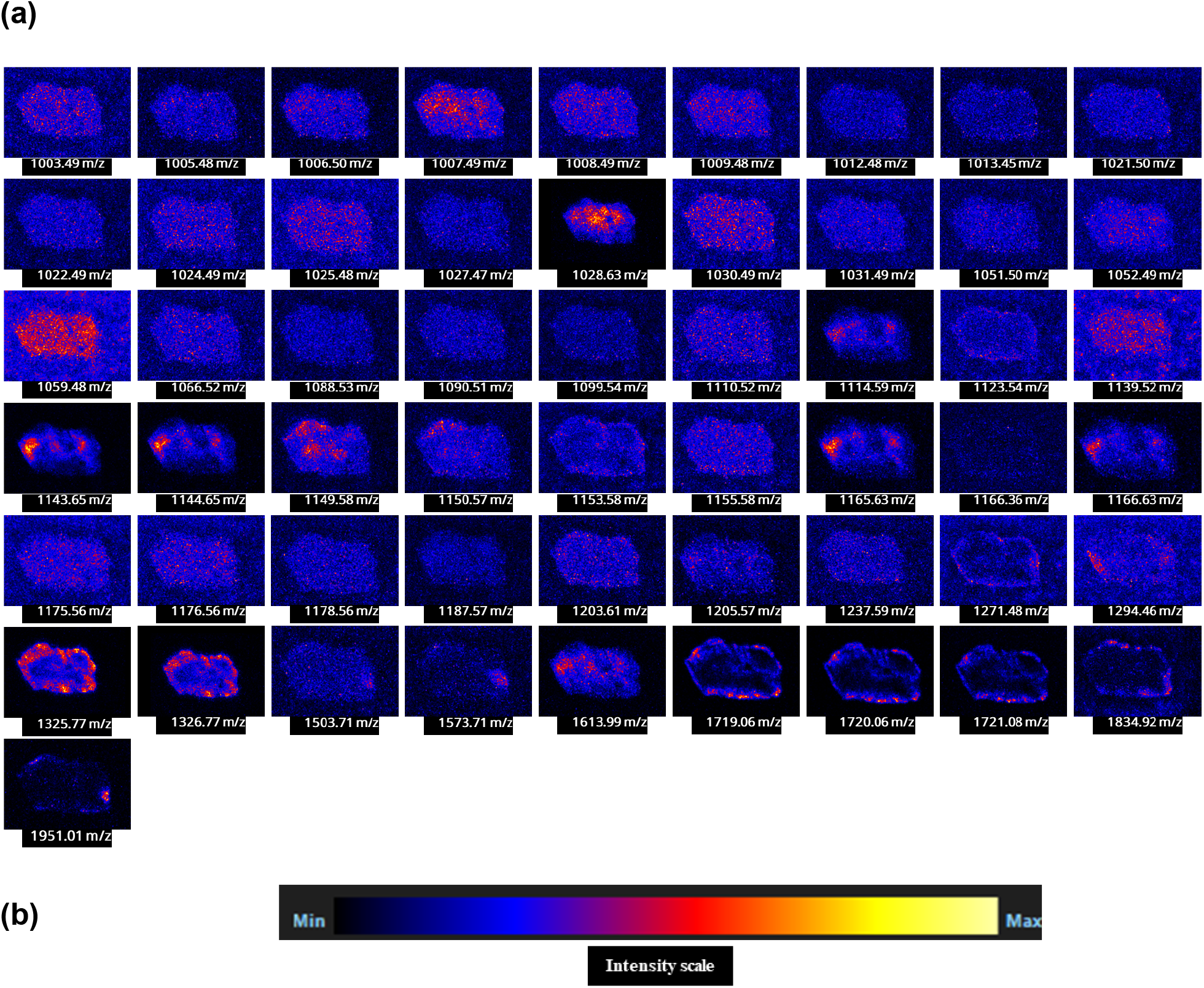
(a) The molecular images of different peptides as represented by their respective m/z values in sample B. The images exhibit hotspots (yellow) within the tissue representing the relatively high intensity of the respective peptide in those regions (b) Intensity scale of the heatmaps where black represents the minimum relative intensity followed by blue, red and yellow representing the highest relative intensity of an ion across the tissue.

**Figure 4:**
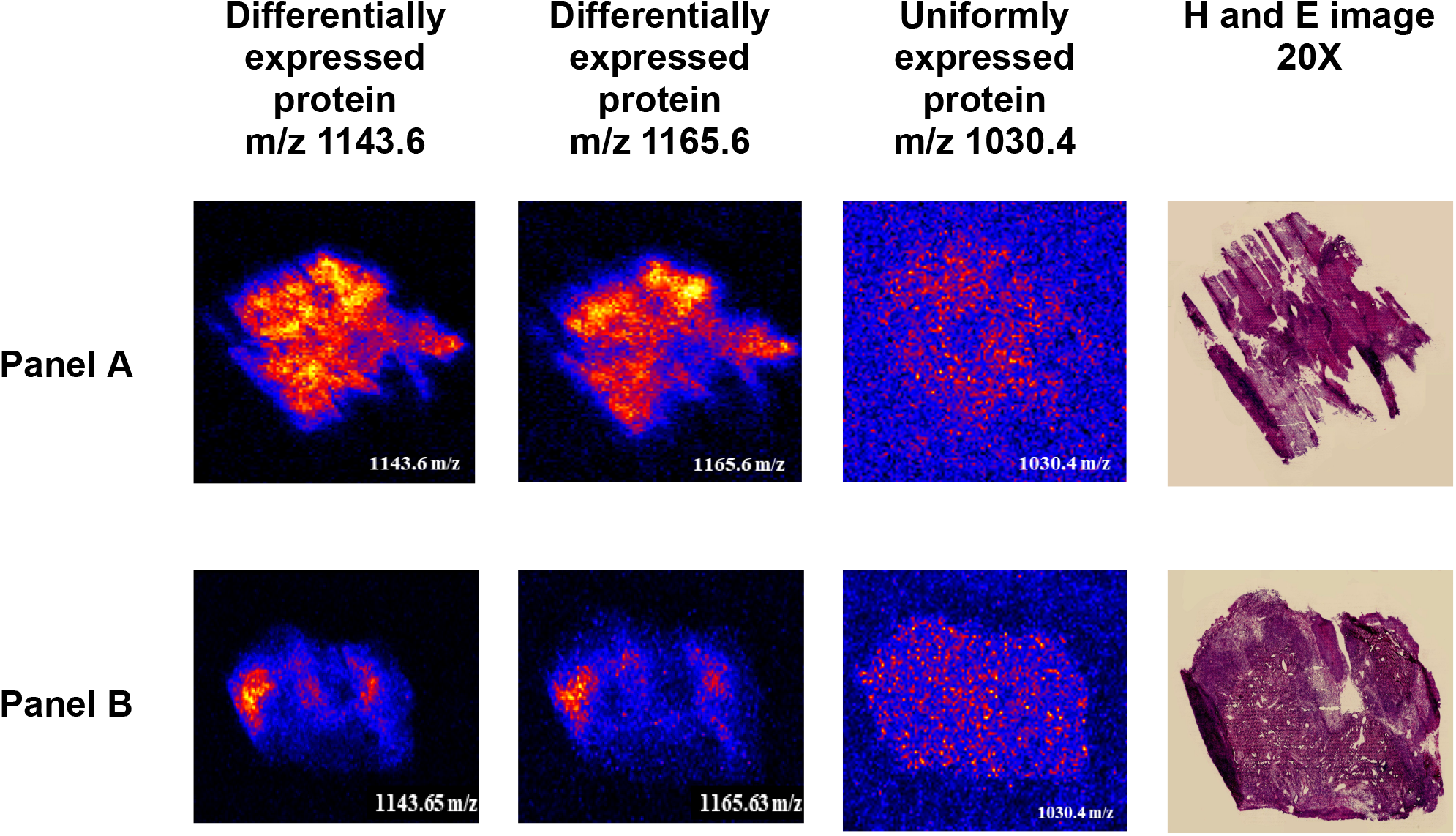
The molecular images of the two representative cervical tissues. Panel A represents the molecular images of sample A, whereas panel B represents the molecular images of sample B. Peptides 1143.6 m/z and 1165.6 m/z are expressed differentially across the tissue sections (red/yellow hotspots) whereas the peptide 1030.4 m/z is uniformly expressed across both the tissues. H and E images of respective tissue sections are shown at 20X resolution.

**Figure 5:**
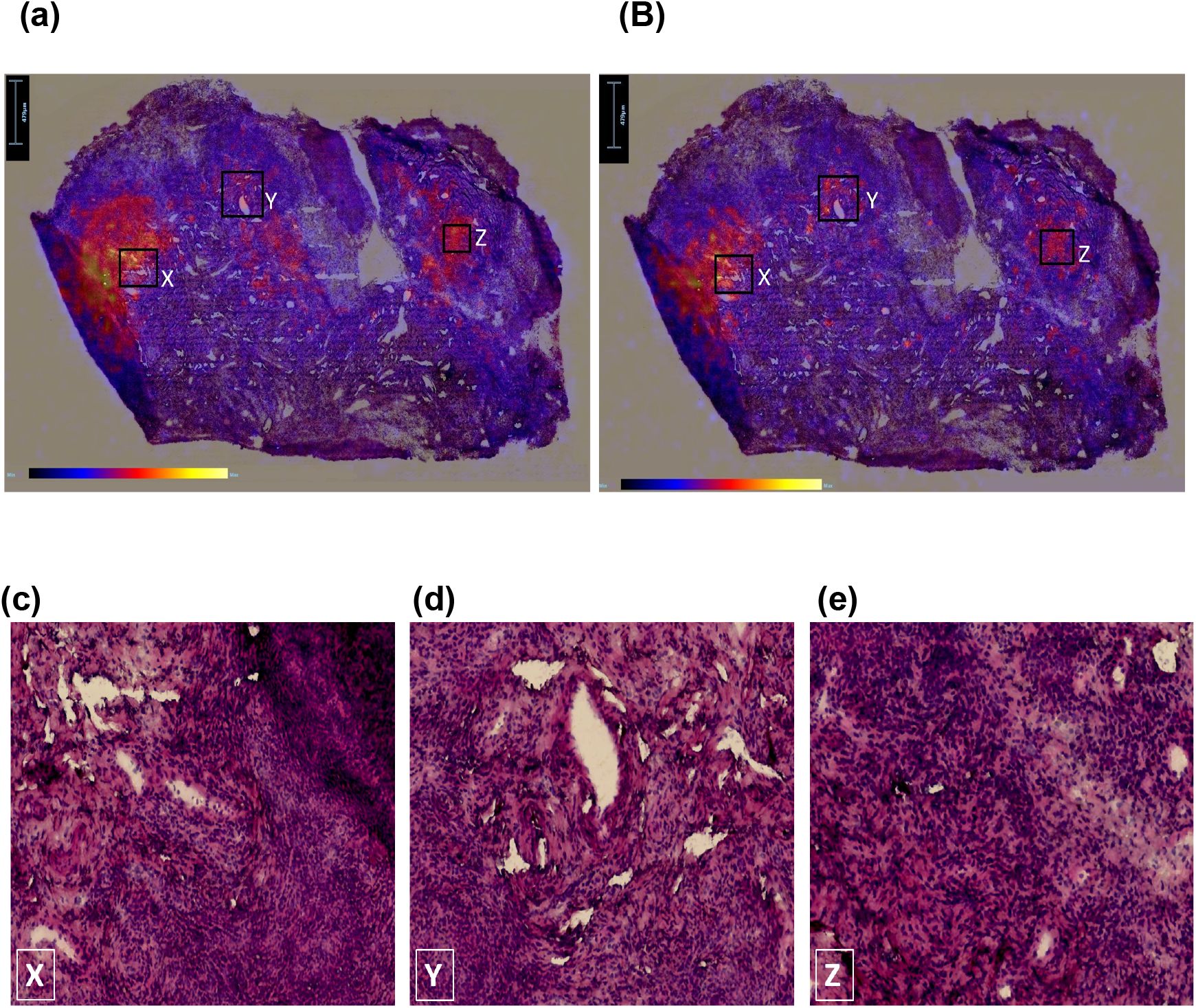
Overlaid images of 1143.6 m/z (a) and 1165.6 m/z (b) with the histopathological image (H and E stained) of sample B. (c, d, e) are enlarged images of the respective regions marked in the overlaid images.

KRT5 is a cytoskeletal protein, generally expressed by squamous epithelial cells and is mainly involved in keratinization [24]. It is generally a marker for less differentiated cells, which are often linked to aggressive tumor phenotypes [25]. KRT5 shows enhanced expression of both RNA and proteins, in several cervical cancer cell lines [24]. It has been associated with aggressive epithelial states in basal like breast cancer, bladder cancer and pulmonary adenocarcinoma [26-29]. It has also been previously reported to be overexpressed causing an increased risk of disease relapse and chemotherapy resistance in ovarian carcinomas [30]. In certain diseases like amyloidosis, it can exhibit aberrant expression and interact with the ECM proteins, however, no such report has been shown till date for cervical cancers [31]. Our study, for the first time, shows an increased localized overexpression of KRT5 in cervical cancer. Prelamin A/C is a protein present in nuclear lamina, and is known to exhibit incorrect cytoplasmic localization in different cancer cells like prostate cancer [32]. Its dysregulation is often associated with genomic instability, aggressive cancer and poor prognosis. It can be over or under-expressed in many different cancer types without a definite overall expression pattern within cancer types and can be good or poor prognosis markers depending on cancer subtype [33,34]. However, the expression profiles of both these proteins in cervical cancer is still largely unknown. Here, with help of spatial proteomics profiling, we found that these proteins show overexpression during the cancer development and progression.

## CONCLUSION

In the present study, we observed differential overexpression of KRT5 and Prelamin A/C in regions marked as cancerous in the cervical cancer tissues. To validate our observations, a larger sample set has to be studied in order to establish these proteins as potential biomarkers of cervical cancer. In addition to biomarker identification, combining label free proteomics with MALDI-IMS can also provide insights into the underlying molecular mechanisms driving carcinogenesis and understand the molecular pathogenesis of these diseases to detect which CIN are more probable to convert into cancers as opposed to benign lesions [37]. This might pave the way for novel diagnostic tools,therapeutic targets and enforcing personalized treatment strategies in cervical cancer.

## ACKNOWLEDGEMENT

We acknowledge all the study participants for providing tissue samples for the study. We acknowledge Dr. Jayasri Das Sharma for her histopathology facility. We also acknowledge the SAIF facility of IIT-Bombay for processing the proteomics samples. Aastha Baliyan acknowledges the fellowship support provided by the CSIR (Council of Scientific and Industrial Research), Government of India to conduct this research.

